# Fast and sensitive multiplexed diagnostic system enabled by real-time solid-phase PCR assay

**DOI:** 10.1101/2025.07.19.665604

**Authors:** Islam Seder, Rodrigo Coronel Téllez, Jing Zhang, Tao Zheng, Stephen McCleary, Saïd El Mouatassim, Jean-François Brugère, Yi Sun

## Abstract

Simultaneous identification of multiple nucleic acid targets is pivotal for high-throughput analysis in clinical diagnostics. Unfortunately, conventional PCR offers limited multiplexing capability due to issues like primer interference, fluorescence spectral overlaps and instrumentation complexity. Solid-phase PCR (SP-PCR), in which primers are physically separated on a solid support, has emerged as an alternative strategy for detection of multiple targets in parallel. However, SP-PCR has been suffering from low efficiency, long reaction time, inability for real-time signal monitoring and accurate quantification, which greatly restricts their applications in multiplexed assays. In this study, we presented for the first time a compact and portable flow-through SP-PCR platform to permit rapid, highly efficient and quantitative solid-phase amplification. We designed an integrated platform comprising a novel mechanically actuated valving system and a dual-chamber SP-PCR system to enable automated sample purification and amplification. Active oscillating the PCR solution between the chambers significantly enhanced the mass diffusion over the solid-phase array. Moreover, after each amplification cycle, the PCR solution was separated from the solid-phase-bound probes, thereby eliminating background signal interference and allowing for real-time monitoring of SP-PCR signals without the need for complex optical systems. The flow-through SP-PCR system demonstrated quantitative detection of five viral pathogens in a single reaction, with a limit of detection of 10 copies per reaction within 20 minutes. This platform provides a promising high-throughput, low-cost and simple-instrumentation multiplexed diagnostic system.

## INTRODUCTION

Polymerase chain reaction (PCR) has been used to detect a few pathogens in one assay to save time and effort in clinical practice (*1*–*3*). For multiplexing, two or more primer sets are designed to amplify different targets, and the detection is commonly achieved via fluorescently labeled probes (*4, 5*). However, multiplexed qPCR instruments require the same number of filters as the number of pathogens, hence limits the multiplexing capacity (*6*). In a typical qPCR instrument, less than five emission channels are usually used, limiting the degree of multiplexing to about 5 due to spectral overlap of fluorophores (*7*). Biofire FilmArray (BFR) multiplex instrument is a notable PCR system that runs multiplexed assay for higher number of pathogens (>15) using a single reading system through compartmentalization of each target in a separate space to target each pathogen separately. However, the BFR system does not offer accurate quantification, and it requires a relatively huge volume for massive compartmentalization.

Solid-phase PCR (SP-PCR), wherein several pathogen-specific probes or primers are immobilized on a solid surface as an array, has been suggested to enable multiplexed amplification and minimize the signal interference in multiplexed detection (*8, 9*). SP-PCR dramatically enhances multiplexing capabilities due to a spatially encoded array that enables targeting multiple specific analytes using a single fluorophore (*10, 11*). However, as SP-PCR is performed in a stationary/static condition, interaction and binding between the generated amplicons of the target genes (in the liquid phase) and the attached pathogen-specific probe (on the solid support) is limited (*12*), which reduces PCR efficiency. Moreover, the signal reading of the SP-PCR process is usually at the end of the assay, and real-time detection is unattainable unless complicated and advanced optical setups are used such as confocal scanning (*13*) and micro-ring optical resonators (*14*).

In this study, we presented for the first time, an integrated microfluidic platform for rapid, highly efficient and real-time SP-PCR assay. The platform incorporated a dual-chamber for SP-PCR, wherein pathogen-specific probes were immobilized on an array in an annealing chamber, and PCR reagent passed back and forth among the annealing chamber and a denaturing chamber. A new mechanical valve was proposed for precise (<0.1 μL variation), bidirectional flow of the PCR reagents. The valve offered robust fluidic control through a structurally simple, single-part design. The approach allowed for the temporary eviction of one chamber, so that the signals of the amplified products on the array could be recorded after each cycle. Moreover, active liquid transportation greatly enhanced the efficiency of SP-PCR. In addition, the two chambers were maintained at constant temperatures, eliminating the need for conventional thermal cycling and significantly reducing transition and ramping time. Furthermore, sample preparation including nucleic acid purification was seamlessly integrated into the fluidic workflow, enabling a complete sample-to-answer diagnostic process. This platform allowed for highly multiplexed real-time quantification with low limit of detection of 10 copies/reaction and rapid turnaround time of < 21 min, while maintaining low-cost consumables and minimal instrumentation requirements. The simplicity and precision of the fluidic architecture make this platform a compelling solution for point-of-care and high-throughput molecular diagnostics.

## RESULTS

### System layout of real-time multiplex PCR assay

The conventional clinic practice for molecular-based pathogen detection is a lengthy, laborious and multi-step process that requires skillful staff and expensive instrumentation (Fig. 1A). Increasing the number of targets per assay, directly increases the sample preparation and instrumentation complexities (*15*). In quantitative PCR (qPCR), the signal measurement instrument is unable to meet with the multiplexing capabilities of reading multiple fluorescent signals due to spectral overlaps of the existing fluorophores (Fig. 1A). Here we designed an automated sample-to-answer SP-PCR system to perform multiplexed assay and enable real-time monitoring of fluorescent signals from the solid-phase array (Fig. 1B and fig. S1). The total system consisted of sample preparation, amplification sections, heating and optical system. Immobilizing multiple specific probes on a solid surface enabled the location-based identification of multiple targets using a single fluorescence wavelength.

**Fig. 1.**
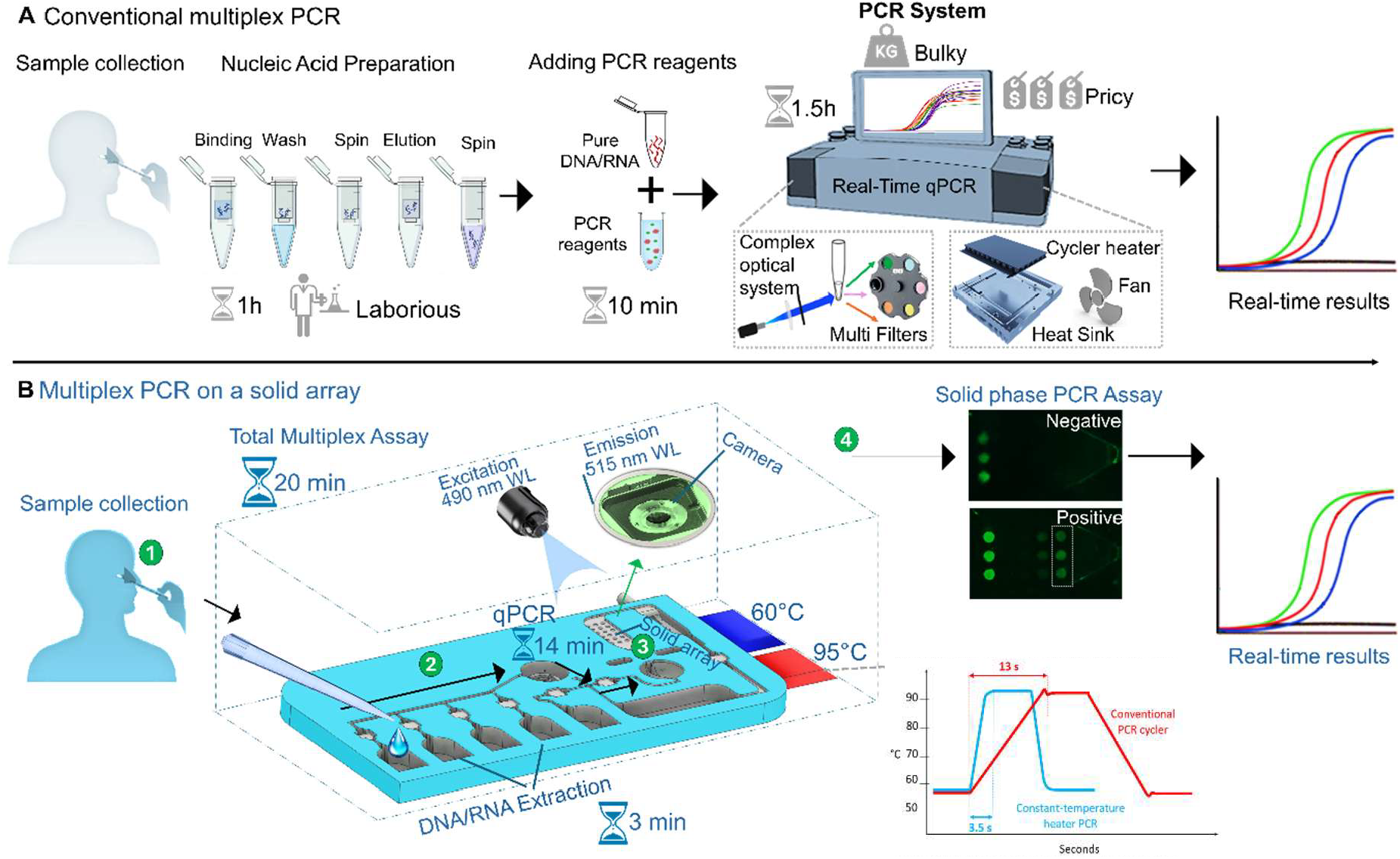
Multiplexed real-time PCR assay on a solid array. **(A and B)** Workflow illustration of conventional multiplex assay (top) and our sample-to-answer multiplexed PCR on a solid array (bottom). The total system contains the nucleic acid extraction and amplification sections on a solid array as well as two heaters and the optical system. In step 1, user loads the sample patient into the chip. In step 2, the system purifies the RNA within 3 min in the extraction section. In step 3 and 4, RT-PCR through solid phase is processed on an array of different pathogen targets with real-time monitoring.

### Fluid control and valving system

Fig. 2A shows the design of the platform. To enable timely and precise transfer reagents for nucleic acid purification and SP-PCR assays, the system integrated a new type of built-in mechanical valving system to control the fluid motion (Fig. 2B). The valve was a single-part rigid cube inserted in a predesigned empty space of a flexible layer. The rigid cube had a channel (0.3×0.3 mm^2^) in the middle of its core. In the closed status of the valve, the channel inside the rigid cube was not aligned with the channel networks of the system, thus fluids were retained in these reservoirs. To open the valve and enable fluid transfer, the valve body was vertically displaced on the z-axis by applying an external force. Alignment of the valve channel with the system channel permitted fluid flow driven by negative pressure (Movie S1). Vertical displacement of the valve was possible due to the elastic property in the layers of the platform. To close the valve, the pressing force was removed to let the valve return to the normal closed status. To prevent undesired fluid flow around the valve wall in the closed status, the size of rigid valve was made bigger than the housing space in the system. We tested several rigid valve sizes with a fixed host space size (2.3 mm^3^ of to the host space in the top flexible layer) and measured the fluid pressure (*P*) inside the microchannel that the valve system can withstand prior to fluid leakage or burst (Fig. 2D). For example, for a 2.3 mm-side cubic space in the flexible layer, a 2.53 mm-side cube valve, which is 10 % bigger than the host space, shows that the fluid bursts at around 40 kPa (Fig. 2D). Such pressure was almost 20 times bigger than the operational pressure of the fluidic system (hydrodynamic pressure drop of the system). This size difference allowed the valve to fit tightly between the flexible layers, forming a sealing mechanism for the valve, and prevents fluid leakage for the undesired reagent from other chambers. For automated pressing of multiple valves (valve 1 to valve 7), a preprogrammed motion of 7 pressing pins was used with the aid of a CAM motion system (fig. S1A).

**Fig. 2.**
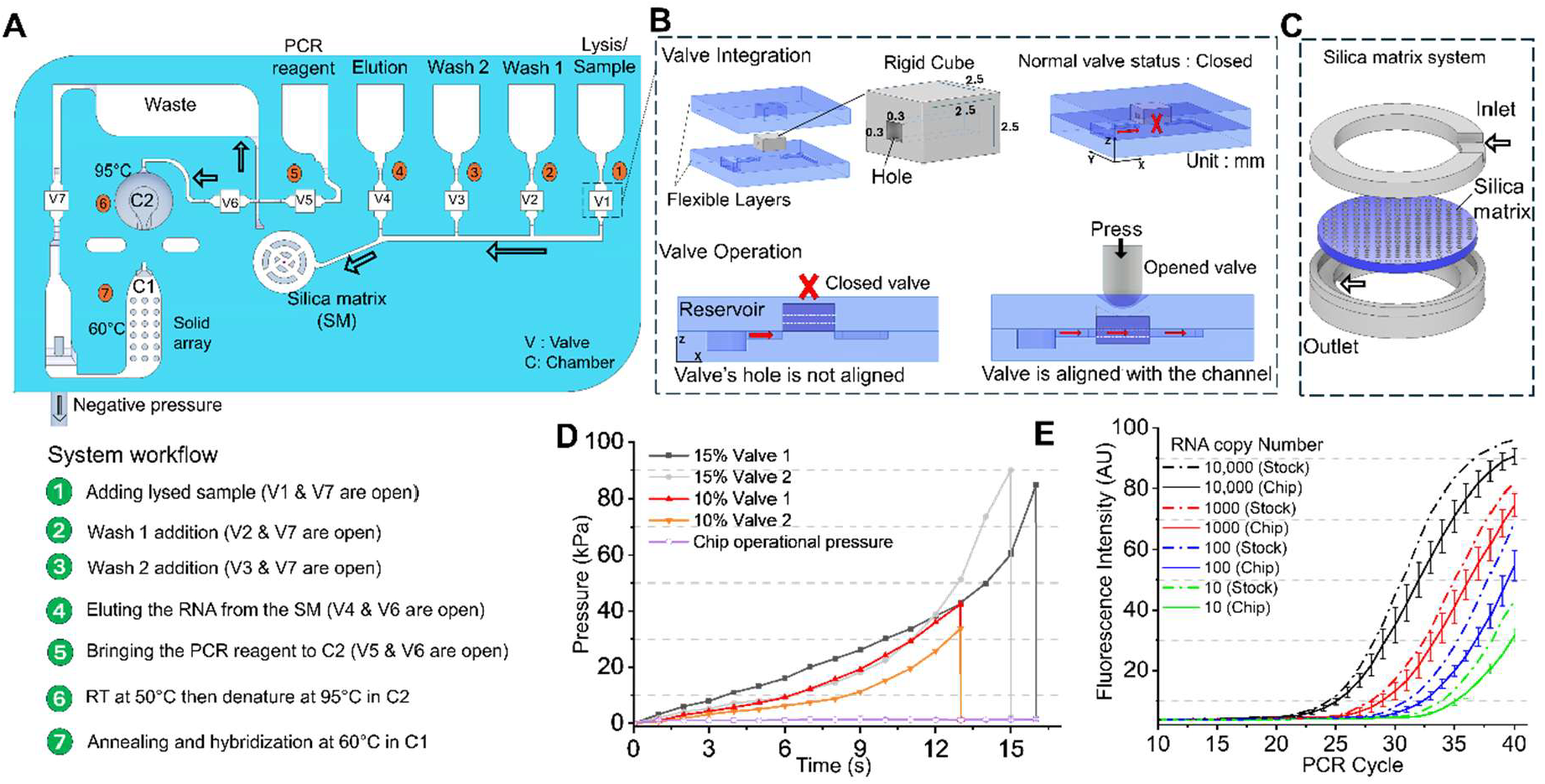
Sample-to-answer multiplex RT-qPCR assay. (**A)** Total assay on the chip. **(B)** The built-in valve system. The rigid body of the valve was inserted in an empty space of the flexible top layer. The fluid was not allowed to pass at a normally closed status of the valve, wherein the channel in the valve core was not aligned with the main channel of the system. To allow fluid motion, the valve was pressed externally, creating a vertical displacement of the valve, hence aligning the valve channel with the system channel network. Rigid valve comes to its original position, closed status, after removing the pressing force. **(C)** Structure of the integrated silica matrix. A top inlet enabled the extraction buffers and the RNA to pass through the silica matrix to capture, purify and release the RNA. **(D)** The valve performance. Pressure burst (maximum pressure that the valve can hold before leaking) vs the chip operation pressure for different sizes of the rigid valve. **(E)** Evaluation of on-chip extracted RNA using benchtop real-time PCR machine.

### Sample preparation

Nucleic acid (RNA) was extracted in the purification section prior to the SP-PCR. In brief, the viral spiked sample was added to the lysis buffer. The lysed RNA was permitted (controlled by valve 1) to pass through the silica matrix (SM), wherein the RNA was captured (Fig. 2A and C). Next, a washing buffer flew through the SM (controlled by valves 2 and 3). Then, an elution buffer was flown through the SM to elute the RNA from the SM and transfer it into chamber 2 (C2) (controlled by valves 4 and 6). The eluted RNA was ready for SP-PCR in the amplification section. We evaluated the performance of on-chip RNA extraction by a benchtop real-time PCR machine to estimate the PCR efficiency (via standard curve) (Fig. 2E). Serial dilutions of RNA samples (10K,1K, 100 and 10 copies) were spiked in the lysis buffer and the purified RNA was collected from C2, then RT-PCR was run for original RNA dilution and the collected RNA, simultaneously (Fig. 2E). Fig. 2E shows a delay in the PCR cycles for the on-chip extracted RNA compared to original RNA stock, indicating an RNA copy loss during the extraction. With an average PCR cycle delay of 0.72, the average purification efficiency was estimated to be 61%±7%.

### Real-time, quantitative SP-PCR strategy

Prior to SP-PCR, RT-PCR reaction mixture was brought from PCR reagent reservoir and mixed with the pure RNA in C2. A reverse transcription step (RT) was performed in C2 at 50°C for 4 min to form complementary DNA (cDNA). Subsequently, PCR was carried out by alternating the mixture between chamber 1 (C1) and C2 during each cycle (Fig. 3A-C). The two chambers (C1 and C2) sit above two temperature-heating zones, 60 °C and 95 °C, respectively (fig. S1A). To move the PCR mixture back and forth between the C1 and C2, an external pin pressed the top side of C2, which is made from flexible material (Fig. 3A). The flexible top side of C2 enabled elastic deformation, thus forcing the fluid to pass to C1 via a connecting channel (Fig. 3A and Movie S1). To bring the fluid from C1 to C2, the pressing force was removed from the top of C2, bringing the deformed C2 to its original volume, thus creating a negative pressure in C2 that forces the fluid to refill C2. The pressing instantly displaced the fluid from C2 to C1 in less than 0.5 s. The temperature of the PCR mixture was rapidly adjusted to the new chamber temperature in a volume dependent manner, *i*.*e*. 3.5 s, 5 s and 10 s for PCR volumes of 10 μl, 15 μl and 30 μl, respectively (fig. S2A). The system exploited the transfer of PCR mixture among the chambers to quickly heat up and cool down the mixture, thus eliminating the need for a bulky thermal cycler (fig. S2B).

**Fig. 3.**
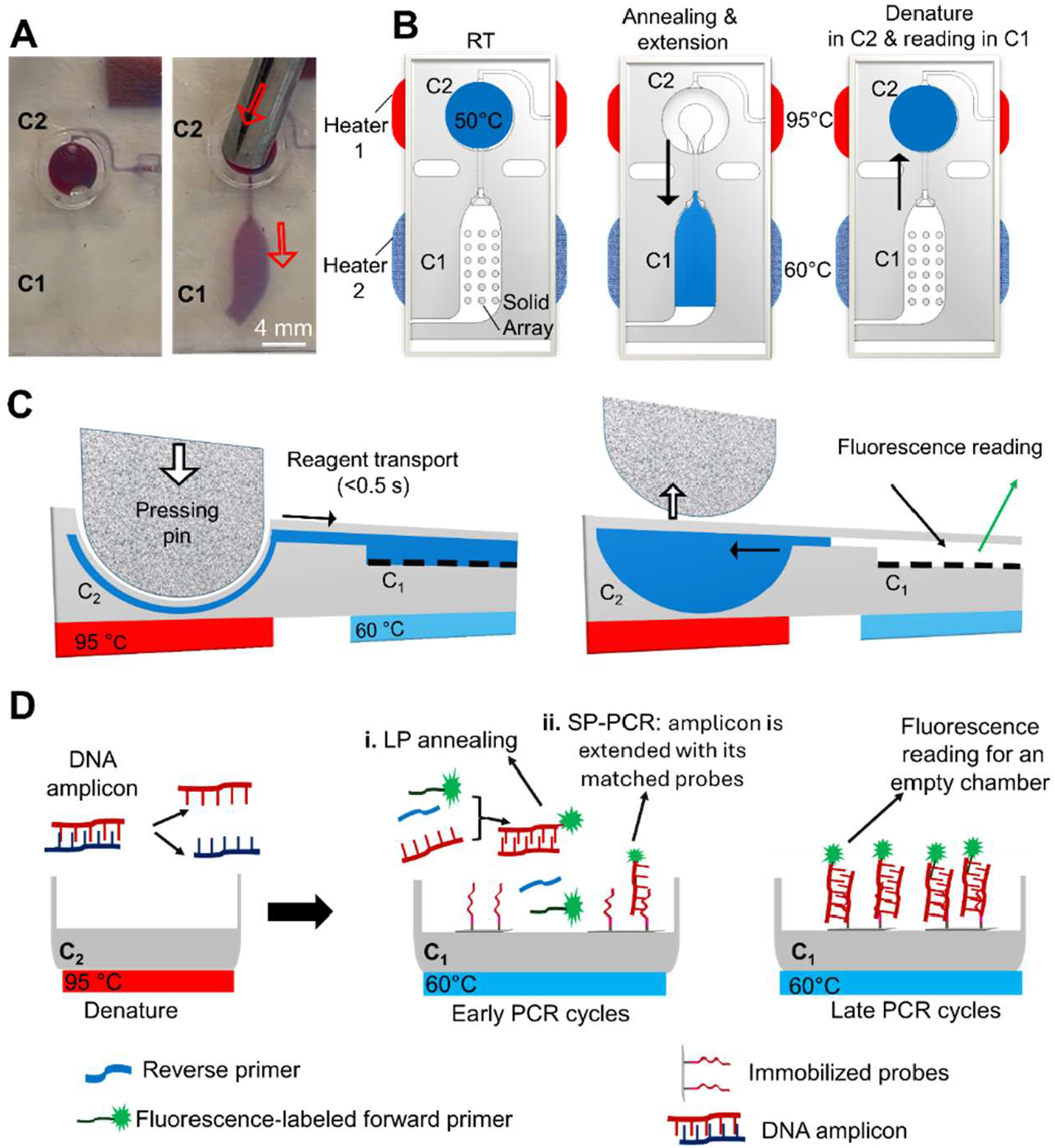
System characterization and solid-phase PCR concept. **(A-C)** Mechanism of fluid transportation during PCR cycling. An external press of the C2 forces the fluid to pass into C1. Releasing the force creates negative pressure in C2, hence quickly retreats the fluid into C2. **(B)** The RT-PCR in two-chamber system. RT-PCR starts with reverse transcriptase (RT) process in C2 to obtain cDNA from the RNA and then PCR process takes place in C1 and C2. **(C)** the side view for the liquid transport mechanism via external pressing pin. **(D)** Mechanism of the solid-phase PCR. The double-stranded cDNA is heated to 95°C to separate it into single strands in C1. The single-stranded DNA pieces (amplicons), along with other PCR ingredients, are in the liquid layer above specific DNA probes that are immobilized onto a solid surface. As PCR mixture is transferred to C2 (60°C), a new DNA is made (via annealing) in the liquid while some of it binds (via hybridization) to the matching probes on the solid surface. As the PCR cycles continue, specific FAM-labeled forward primers extend to the appropriate sites on the template. More DNA is produced, which also acts as a new template for further amplification on the solid surface.

For each PCR cycle, the cDNA was denatured at C2, then the reaction mixture was transferred to C1 for the annealing and extension step (Fig. 3B and 3C). The annealing and extension occurred in both liquid phase and solid phase. The forward primer was fluorescently labelled with FAM. During the annealing step, new amplicons were formed in the liquid phase with forward and reverse primers annealed to the denatured strands and got extended, while portions of the denatured strands (with FAM labeled forward primer) were hybridized to the immobilized probes on the solid-phase array (Fig. 3D). The probes were elongated and the FAM labeled strands stay on the solid-phase assay to provide the fluorescence signal. Such primer labeling sequence strategy enabled detection using a single spectral range (single emission wavelength).

After each PCR cycle, the PCR mixture that contained FAM-labeled forward prime and reverse primer was transferred to C2 to eliminate any source of signal interfering with the array (Fig. 3B), allowing for real-time signal reading of the solid array. Thus, the fluid reciprocated among the two chambers to i) change the temperature of PCR mixture from 60 °C (in C1) to ∼95 °C (in C2) or vice versa, ii) temporally empty C1 for real-time fluorescence signal reading of the solid array after each PCR cycle while denaturation of liquid amplicons is going on in C2; and iii) provide a continuous and dynamic motion for the PCR solution.

### Performance of two-chamber SP-PCR

Initially, to evaluate the proposed dual-chamber PCR system, a traditional liquid-PCR test was performed for a single target (Covid-19 virus) with SYBR green dye. Fluorescent signals were intensified as the PCR number increased, proving the functionality of the system (fig. S2C, fig. S3A). Then, we evaluated our dual-chamber SP-PCR concept via immobilizing 3 spots of specific probe of Covid-19 virus, 3 spots for negative control (oligonucleotides having no homology to the target sequence) and 3 spots for FAM probe on surface of C1 (Fig. 4a). RNA and RT-PCR reagents were added in C2. Then, SP-PCR was performed through oscillation of the assay mixture between C1 and C2. Fluorescence signal of the solid panel was plotted in real-time after each annealing step in C1 to determine the assay performance (Fig. 4B). Fluorescence measured on the microarray spot increased linearly with the concentration of amplicons in solution and on the solid spots. Since all the PCR reagents are dragged away from the reading chamber (C1), no wash steps are required to reduce fluorescent background. Fig 4B indicates that the signals of the FAM probe remained relatively linear over the 40 PCR cycles indicating the stability of the chemical bonding through a 40 times fluid alteration among C1 and C2. Meanwhile the specific spot for Covid-19 displayed an increase of fluorescent signal as the PCR cycle increased due to the solid-phase PCR reaction. The sensitivity (limit of detection) of the assay was 10 copies/reaction with PCR efficiency of 83.4% for single target of covid-19 virus (Fig. 4C).

**Fig. 4.**
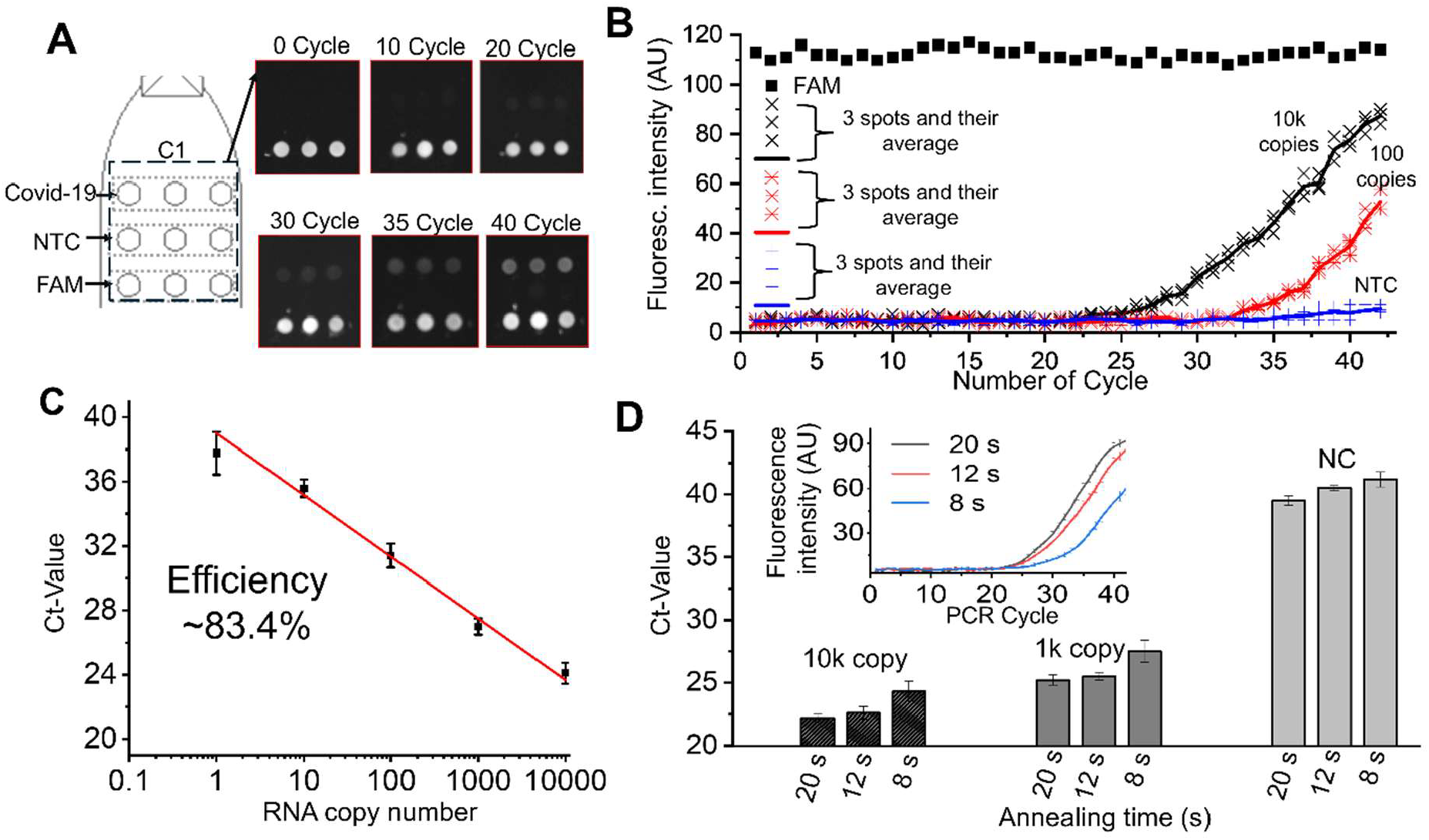
Characterization of dual-chamber SP-PCR. **(A)** Real images of the solid-phase PCR for a single target (Covid-19 virus) in C1. The immobilized FAM shows stable and strong signal through the PCR cycle increment, while the signals of target spot (Covid-19) gradually increase with the increment of the PCR cycles. **(B)** Real-time signal reading for the array spots for Covid-19 virus with copy number of 10k, 1k and non-template target (NTC). The line represents the average of the 3 measurements of the 3 spots of the array. Annealing and denature time are 20 s and 5 s, respectively. (**C**) The standard curve for PCR efficiency calculation for the SP-PCR with a single target (Covid-19 virus). **(D)** Influence of annealing time (hybridization) in C1 on PCR performance. The insert of graph (D) shows the real-time fluorescence with PCR cycles.

The displacement of the PCR reagents from C2 to C1 occurred in less than 0.5 s, creating a 2000 μL/min fluid velocity. With such velocity, a turbulence flow was created inside the C1 (fig. S4 and Movie S2). The suggested dual-chamber platform therefore generated an additional fluid motion i.*e*. advection, due to the transporting of PCR solution. The dynamic motion of PCR components enhanced the diffusion process between the PCR solution and the solid array/immobilized probes, which resulted in a high SP-PCR efficiency.

To reduce the assay turnaround time, we also optimized the annealing/extension time (Fig. 4D). Apparently, a minimum of 12 s is required for the assay without compromising the assay efficiency for a ∼150-nt specific extension, thus unless mentioned otherwise, we used 12 s annealing/extension for SP-PCR assay. Such annealing time was important to reduce the essay turnaround time.

### Sample-to-answer multiplexed real-time detection

To evaluate the integrated system for multiplexed real-time assay, we designed an array of specific probes for 5 viral pathogen targets (Covid-19, Influenza A (Inf. A), Influenza B (Inf. B), Rhinovirus and Parainfluenza (Para-inf.) viruses) (Fig. 5A). RT-PCR was performed in the two-chamber PCR section for the 5 viral targets after the RNA purification in the extraction section. Figure 5B depicts the fluorescence singles of the solid array in real time after the annealing/hybridization process. The assay successfully distinguished the existence of 5 viruses for the 5 positive cases (spiked viral RNA in lysis buffer). Figure 5C shows 3 positive tests in 3 lanes for the array for the three spiked viral RNA (Covid-19, Inf. A and Rhino viruses) while two lanes show no signal, since no viral RNA was spiked (no template control (NTC)). The average LOD for the 5 viruses was 10 copies/reaction. The proportion of negative test results out of truly negative samples, i.e. specificity, was 100% for the cutoff at cycle 38 (Fig. 5E-I). Slightly progressive signals started to build up on the solid spot after the 40 cycles that limit accuracy to distinguish between the positive from negative sample. Hence, we considered a signal after 38 cycles as a negative sample, as running the PCR above 40 cycles showed a slightly stronger signal for samples with low copy numbers (<10 copies) than the negative control (NTC) (fig. S5).

**Fig. 5.**
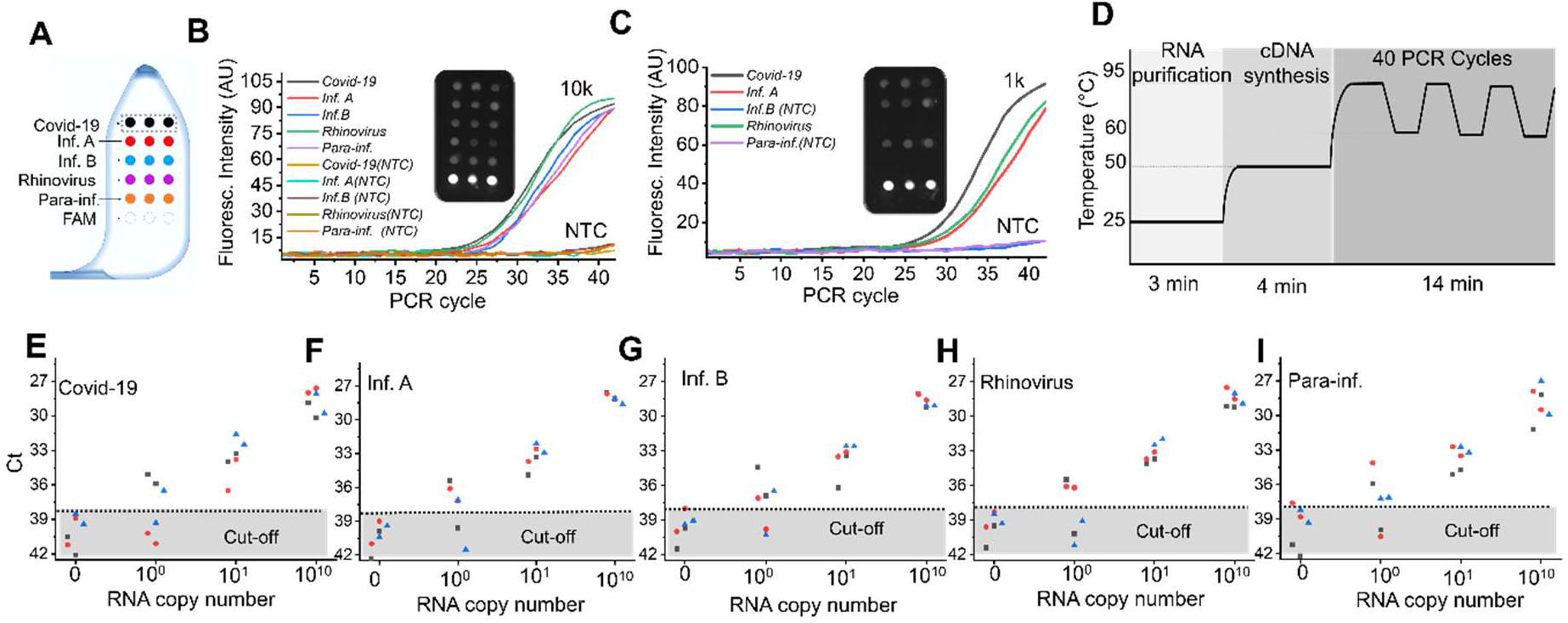
Multiplexing via solid array assay. **(A)** Arrangement of the immobilized probe in C1. **(B)** Multiplexed PCR for 5 viral pathogens run simultaneously with 10,000 RNA copies and 0 copies. **(C)** Multiplexed PCR for 5 viral pathogens with 3 positive lanes (3 viral RNA was spiked) with 5 lanes containing probes immobilized for 5 viral RNA). **(D)** Assay total time. **(E-I)** LOD for Covid-19, Inf. A, Inf. B, Rhinovirus and Para-inf., respectively.

With instant transporting and the rapid heating/cooling of the PCR mixture, a short turnaround time of the multiplex assay is achieved. The sample-to-answer test takes ∼20 min (Fig. 5D) to perform multiplex assay, including 3 min for RNA extraction, 4 min RT process and 14 min for 40 cycle PCR of 15 μL volume.

## DISCUSSION

We have demonstrated a real-time sample-to-answer assay based on solid-phase PCR array for multiplexed detection. The proposed system allowed real-time signal reading, hence quantification, for solid-phase PCR without using external or special optical equipment. The real-time quantification was enabled through moving the PCR solution away from the solid-phase array after each annealing step, hence reading the signals of the array in an empty space. To allow the fluid movement, a two-chamber PCR system was integrated, where DNA annealing, extension and signal reading occurs in one chamber (C1) and DNA denature occurs in another chamber (C2). To achieve the sample-to-answer assay, that includes several reagent additions, we have also introduced a new built-in valve system enabling precise and simple reagent manipulation for both nucleic acid (RNA/DNA) purification and amplification steps. To assess the utility of the total platform in clinical scenarios, we performed multiplexed viral detection for 5 viruses simultaneously, analyzed the impact of PCR dynamic reagent motion on solid-phase PCR assay sensitivity, and characterized the valve system properties of manipulating several assay reagents and buffers.

For manipulating the assay reagent and performing nucleic acid purification, the valving system is an essential component for sophisticated assay performance. To ensure fluid control robustness, the conventional valves often include several parts that increase the system integration complexity, whereas the passive valving, that usually depends on the surface tension of the chip material and storage condition, lacks robust performance. Hence, we have developed a built-in single-part valve for precise fluid control. The valve consisting of a rigid cube with a hole inserted into a predesigned flexible space, trapped buffers in reservoirs as long as the central hole remained misaligned with the chip channels. When externally pressed, the valve aligned with system channels, enabling fluid flow. The elastic platform layers facilitate vertical displacement, ensuring an instant response and preventing inner fluid leakage. The prevention leakage mechanism was guaranteed by designing the rigid cube slightly larger than its housing space, creating a tight seal. The mechanical motion provided an instant and precise response for fluid control. In comparison to several valving systems in the literature that involve a complex level of system integration, such as piezoelectric (*16*), electrostatic (*17*), and pneumatic valves (*18*), our press-to-activate valve is extremely simple (single part), precise (instant response), and reliable (mechanical actuation), while the valving system does not require additional layers or oil to prevent fluid leakage.

With the integration of valve system (7 valves) and a silica matrix, we designed an auxiliary unit for nucleic acid purification on the system in order to provide sample-to-answer test without intervention (Fig. 3), greatly easing the overall sample testing and avoiding contaminations. The nucleic acid purification took 3 min on the disposable chip with fully automated operation.

After the nucleic acid (RNA) was purified, the sample was transferred to chamber C2. We designed a dual role for the chamber C2, initially devoted to the RT reaction, and then for the denaturation step of each PCR cycle, which simplified the device structure and the workflows. For the SP-PCR, two chambers (C1 and C2) communicate with each other through a narrow channel allowing the PCR reaction mixture to be reciprocated back and forth between the two chambers, with two different heating zones, for PCR thermal cycling. Traditionally, strong fluorescence in PCR solution originated by the freely labeled primers, prevents a real-time reading for solid array unless a special background suppression or high signal-to-noise ratio mechanism are used, such confocal scanning (*13*), surface plasmon resonance spectroscopy (SPR) (*15*), opto-fluidic equipment (*8*), wavelength shift of a silicon micro-ring optical resonator (*14*), and microfluidic spatially defined droplet arrays (*19*). Those alternative techniques involve complicated and excessive optical components, moving objectives for fluorescence scanners or opto-fluidic equipment. Here, we propose the simple two-chamber solution that allows the PCR reagent to be removed away from the solid-phase array in C1, thereby enabling clear fluorescence reading. In C1, real-time signal reading of the amplified PCR products on the immobilized probes was enabled by performing hybridization for labeled amplicons to the complementary probes on array. Because the PCR mixture that contains the unattached FAM-labeled primers was driven away to C2 during each PCR cycle, the signal of the amplicon on solid array was distinguishable without the need for extra optical tools. It is noted that an enhancement of the signal at each PCR cycle could be implemented, for example by using fluorescent dNTPs incorporated during the SP-PCR and allowing more than one fluorescent molecule to be present on each SP extended strand. We believe that enabling real-time monitoring of solid-phase amplification signals represents a significant advancement in SP-PCR technology, and will greatly enhance its usefulness across a wide range of applications where absolute quantification is required.

The two-chamber PCR system also provided rapid temperature changes (Fig. S2B), thereby significantly reducing assay processing time as compared to known solid array systems (*1, 13*). Usually, a relatively big PCR volume (>15 μL) is necessary for reliable SP-PCR assay, since small volume reduces the target copies and increases fluid handling issues such as evaporation or complexity in handling small volume. However, with large PCR volume, the heating up and cooling down of the PCR solution is lengthy (>1h). Besides, PCR cycler requires a bulky thermal cycling system that consists of a heater, a fan, a heatsink and a precise temperature controller. Here, the suggested dual-chamber PCR system enables a fast thermal response, since the PCR solution moves to a preset temperature compartment. Hence, there is no need to heat up and cool down the whole system. The PCR solution is rapidly adjusted to the wall temperature of the compartment. Of course, in our system, the temperature transition time can be further reduced by using smaller volumes and increasing surface-to-volume ratio of C1.

Moreover, the assay performance enhancement in sensitivity and shortened annealing process time, can be attributed to the dynamic motion of the PCR reagent above the solid array. Conventional SP-PCR runs in a static reaction above a two-dimensional solid array, wherein passive diffusion process between the PCR solution and the solid array is the dominant mechanism that enables interaction between the immobilized probes and the PCR components. Pierik et. al. (*13*) only obtained 58 % PCR efficiency after hybridizing amplicons in a static SP-PCR. In contrast, our flow-through SP-PCR achieved more than 80% PCR efficiency, far above the traditional stationary SP-PCR. The results suggested that enhancing the diffusion of PCR products near the solid array surface played an important role in improving the amplification efficiency of the solid-phase PCR

In practical terms, the proposed set-up allows a practitioner to inject the patient sample (nasal swab or saliva) directly into the microfluidic device and obtain the multiplexed assay result in ∼ 20 min without compromising any purification or amplification steps. Comparing to commercialized multiplexed assay platforms, such as Filmarray 2.0 and BioFire® SpotFire® from Biomerieux Inc., that require 45 min and 15 min for the sample-to-answer detection, respectively, our system provides a competitive assay time with a simplified system instrumentation. The dual-chamber PCR system presents a scalable multiplexed assay for several pathogen targets. The assay is fast and sensitive with real-time quantitative detection capabilities. The average LOD of several tested viral pathogens was 10 copies/reaction. Below such copy number, the possibility of a false positive results increased, due to the interaction between the non-specific products and the immobilized probe.

We expect that clinical application of the suggested multiplex real-time detection platform with sample-to-answer capabilities will be both straightforward and cost-effective, with huge potential adoption in clinics or point-of-care areas. The proposed system is applicable for disease screening for poor-resource clinics or national ports for its structure simplicity and rapid turnaround assay time during an epidemic or pandemic pathogen outbreak, with ability to design and implant certain group of targets on solid-phase array. Additionally, the simple thermal system drastically reduces the size and weight of the system PCR, making it a suitable system for drive-throw tests with significant ramping up and cooling down time reduction. We tested only 5 pathogen targets on an array as proof of concept for the multiplexing capabilities. However, printing a larger array is obtainable as we have shown in figure S6. A yet further aspect of providing a solid array on a separate substrate, which can be slotted in the second chamber during the chip production, may improve manufacturing flexibility for targeting the total analysis system to different analytes. This allows the system’s user to utilize the same PCR platform for producing different batches of chips directed at different analytes.

## MATERIAL AND METHODS

### Platform structure and fabrication

The platform consists of a disposable device and permanent auxiliary parts. (fig. S6) The disposable device (30×50mm^2^), wherein the assay takes place, consists of a bottom layer of a 3-dimensional (3D) printed biocompatible material (Biomed, Formlabs) and a top layer of flexible polydimethylsiloxane (PDMS) material. The 3D-printed bottom layer has an engraved space (35×4.5×2 mm^3^) to host a flexible PDMS cuboid that allows structure deformability during the valve operation. Chambers and the connecting microchannels lie in the bottom layer (fig. S1B). The top PDMS layer contains only the cubic grooves for the rigid valve. Flexible material is used for the top layer since the operation of the valves and C2 requires material deformation. PDMS is also biocompatible with PCR process. Because the PDMS is an air permeable material, that increases the evaporation of the PCR solution, bottom and side walls of the disposable device were rigid 3D-printed material. Detail of fabrication and the platform components are listed in Table S1. The valve cube is made of a 3D-printed rigid material. The body of the valve was inserted inside the grooves of the top layer (fig. S1B). The permanent parts of the platform are two constant-temperature heaters fixed at the bottom of the device, excitation and emission filters with a camera fixed at the top of the device, and a CAM system for fluid control. The CAM system consists of 7 CAMs, 7 followers, 7 pins and 7 springs to mechanically control the valve and the fluid motion in C2. A stepper motor is used to control the motion of the CAM (fig. S1 and Movie S3).

### PCR chamber design

The two chambers (C1 and C2) for the PCR process possess different shapes and features since they perform different steps of annealing and denture, respectively. C1 was designed shallow (250μl-depth) and long (3 mm-length) to increase the surface-to-volume ratio, hence increasing the heat dissipation of the hot PCR mixture (95°C) coming from the C2. C2 is designed as a hemisphere shape to enable a complete fluid transfer once the pressing-pin is engaged since the pressing-pin tip has a hemisphere shape. A microchannel (130 μm x 130μm x 2700 μm) connects C1 and C2 for the fluid alternating among the chambers. The microchannel cross-section (A) was small to increase the fluid velocity (v) upon the fluid transporting among the chambers, given that the volumetric fluid flow rate (Q) is proportional to v and A (Q = v × A). The microchannel was long (2.7 mm) to prevent a thermal gradient establishment between C1 and C2 although this length creates a microchannel total liquid volume of ∼0.4μL (2 % of the total assay volume). The microchannel was on the top of C2 hemisphere to allow any generated bubble to pass forward to C1 during the pin pressing while the microchannel has smooth-rounded edges at the C1 side to enable smooth fluid motion from C1 to C2.

### Surface functionalization and probe immobilization

Attachment of the probe to the surface of the array compartment is performed via functionalizing the surface of the microfluidic space. The surface of the C1 was functionalized with aldehyde group (−CHO) prior to the probe immobilization (Fig. S7). Initially, we tried to functionalize the 3D-printed material bottom of C1. However, unrepeatable probe density and uniformity was detected in specific spots (Fig. S8A), thus we coated the 3D-printed surface of C1 with PDMS since PDMS has hydroxyl group (OH-) upon oxygen plasma treatment then proceed for PDMS surface functionalized for the solid array. Immobilization of probes harboring a C6-amino linker to PDMS surfaces was performed as reported (*22*) with some modifications. Briefly, the PDMS layer above the 3-D printed surface was thoroughly cleaned with acetone, RNase-free water, and dried with N_2_. Next, the surface was subjected to oxygen plasma treatment for 5 minutes and immersed in 5% (3-Aminopropyl) triethoxysilane solution. After overnight incubation at room temperature, the surface was washed with RNase-free water, dried with N_2_, and cured for 1 hour at 110°C. Afterward, the surface was immersed in a 5% glutaraldehyde solution for 2 hours at room temperature, washed with RNase-free water, and dried with N_2_. Finally, 3 μL of each probe (100μM) were spotted on their respective lane by employing the column array dispenser developed by our team (fig. S8 and Movie S3). Finally, 1% bovine serum albumin (BSA) in phosphate-buffered saline was added in C1 and C2 to reduce a non-specific binding then washed by RNase-free water. Probe attachment and uniformity on the spots were ensured by allowing the dispensing columns that contain the probe solution to be in continuous contact with the treated PDMS surface for 12 hours at room temperature under controlled humidity conditions (humidity index ∼98%). The stability of the immobilized probe against the continuous motion of the PCR mixture is shown in the unchanged intensity of FAM probe after 40 cycles of fluid transporting among C1 and C2 thanks to the strong chemical immobilization.

### Primer and probe design

Primers and probes designed for this study are presented in Table 1. A FAM fluorophore was included in the 5’-end of forward primers to allow the detection of PCR-amplified products. A C6-amino linker was included in the 5’-end of the probes to facilitate attachment to PDMS surfaces. Forward and reverse oligos were purchased from TAG Copenhagen and probes were purchased from IDT.

### Array arrangement

To track and confirm probe immobilization, a probe with a random nucleotide sequence was designed with a C6-amino linker on its 5’-end and FAM fluorophore on its 3’-end (Immo-P, Table S2). The fluorescent signal of this probe was assessed after probe immobilization and PCR amplification (fig. S8A). Once the immobilization protocol was optimized and the probe attachment ensured, this reporter probe was omitted from the final multiplex RT-PCR array. Both the forward and reverse primers were applied in 0.4 μM final concentration.

### RNA samples

Synthetic RNA controls for Covid-19 (SARS-CoV-2, MN908947.3), Influenza A H1N1 (NC_026433), Influenza B (NC_002209), Rhinovirus 89 (NC_001617.1), and Parainfluenza 1 (NC_003461.1) were purchased from Twist Bioscience.

### Nucleic acid purification and PCR buffers

RNA purification kit (QIAamp Viral TNA Min Kit, QIAGEN) was uploaded in their predesign reservoirs as follows: 50 μl, 50 μl, 50 μl, 10 μl for lysis, wash 1, wash 2 and elution buffers, respectively. For a 20-μL RT-PCR test, the SensiFAST™ Probe No-ROX One-Step kit from Meridian Bioscience was used. 16μl of RT-PCR master mix, that includes 6μl of purified RNA, were combined with 4μl of a solution with all 10 forward and reverse primers mixed in equimolar concentrations (0.4μM). The volume of the reagents is scaled up or down according to the total volume of the assay. RT was performed in C2 at 45°C. Next, the C2 temperature increased to 95°C to allow polymerase activation for 1 min. Unless specified, PCR amplification was performed by promoting cDNA denaturation in C2 for 5 seconds at 95°C and then transferring the solution to C1 to perform annealing and extension for 12 seconds at 60°C.

### Fluorescence measurements

The fluorescence intensities of all spots shown on the array are measured at the end of the annealing step of the liquid-phase PCR (the hybridization step of the solid-phase PCR) using a CMOS camera (CS165MU/M) and ThorCam software. We used the average fluorescent intensity of three spots of the same target, i. e. probe sequence, for each measurement. We normalized the recorded signals to the measured fluorescence intensity of the C6-amino linker-FAM fluorophore. For the traditional liquid phase PCR (Fig. 4B), the fluorescent signal was produced when EvaGreen® dye (expiation at 498nm-wavelength) activated after binding to the double-stranded DNA (dsDNA) generated. The full image for C2 was taken after each cycle, wherein the fluorescent signal was proportional to the dsDNA present in the reaction. Details of fluorescence signal acquisition are in SI.

## Acknowledgements

This work was supported by NovoNordisk Foundation, Exploratory Interdisciplinary Synergy Programme, Grant no. NNF21OC0070706.

## Conflict of Interests

I.S and Y.S. are inventors on a patent concerning the valves described here. J.-F. B and S.MC.L are inventors on the patent PCT/EP2025/061585 describing the solid-phase PCR with moving liquid reaction among C1 and C2.

## Supplementary Materials

The PDF file includes:

Sections S1 to S7

Figs. S1 to S8

Other Supplementary Material for this manuscript includes the following:

Movie S1 to S3

## Notes

### Competing Interest Statement

I.S, R.C.T. and Y.S. are inventors on a patent concerning the valves described here. J.-F. B and S.MC.L are inventors on the patent PCT/EP2025/061585 describing the solid-phase PCR with moving liquid reaction among C1 and C2.

