## Supplementary for "Fast and sensitive multiplexed diagnostic system enabled by real-time solid-phase PCR assay"

### Material selection and fabrication

We used 3D-printed material for the bottom layer of the chip and the valve and polydimethylsiloxane (PDMS) material for the top layer and for the cuboidal groove in the bottom layer. Using two different materials to fabricate a single device is not favorable since it increases the fabrication steps. However, because the chip functionally requires flexile material from the top and air-impermeable material on the bottom, to reduce the evaporation, especially in small PCR volumes (<15µL), we integrate the layers with two different materials. The platform used a solid 3D-printed polymer on the side and bottom walls of the PCR chambers. Flexible PDMS for the top layer is needed for the operation of valve and C1 while PDMS also possesses biocompatibility with PCR process. Details of the fabrication process are found in Figure S2 and S5. The 3D-printed mold was designed by Autodesk software. The PDMS layer was fabricated from a 3D-printed mold. Grooves to host the rigid valve were the only features in the top layer. For 3D-printing process, the parts were soaked in isopropanol for 5 min, sonicated for 5 min, exposed to UV light for 15 min, and left in an oven at 80 °C for 8 h. Then, a PDMS mixture 10:1 (10 Sylgard 184 silicone and 1 Sylgard 184 silicone curing agent) was poured on the mold and left in the oven for PDMS crosslinking at 60 °C for 2 h. Such curing conditions create a Young's modulus (elastic modulus) of 1.5-2.5 MPa and a Shore A hardness of ~40 (***1***). Unlike the top side of the chip, the valve was made from a 3D-printed part to have a rigid body and thus it sustains the mechanical stress during the valve pressing without experience deformability. The chassis box of the system was made of 3D-printed parts to provide a dark environment for the chip during the signal reading and to provide rigidity to the system.

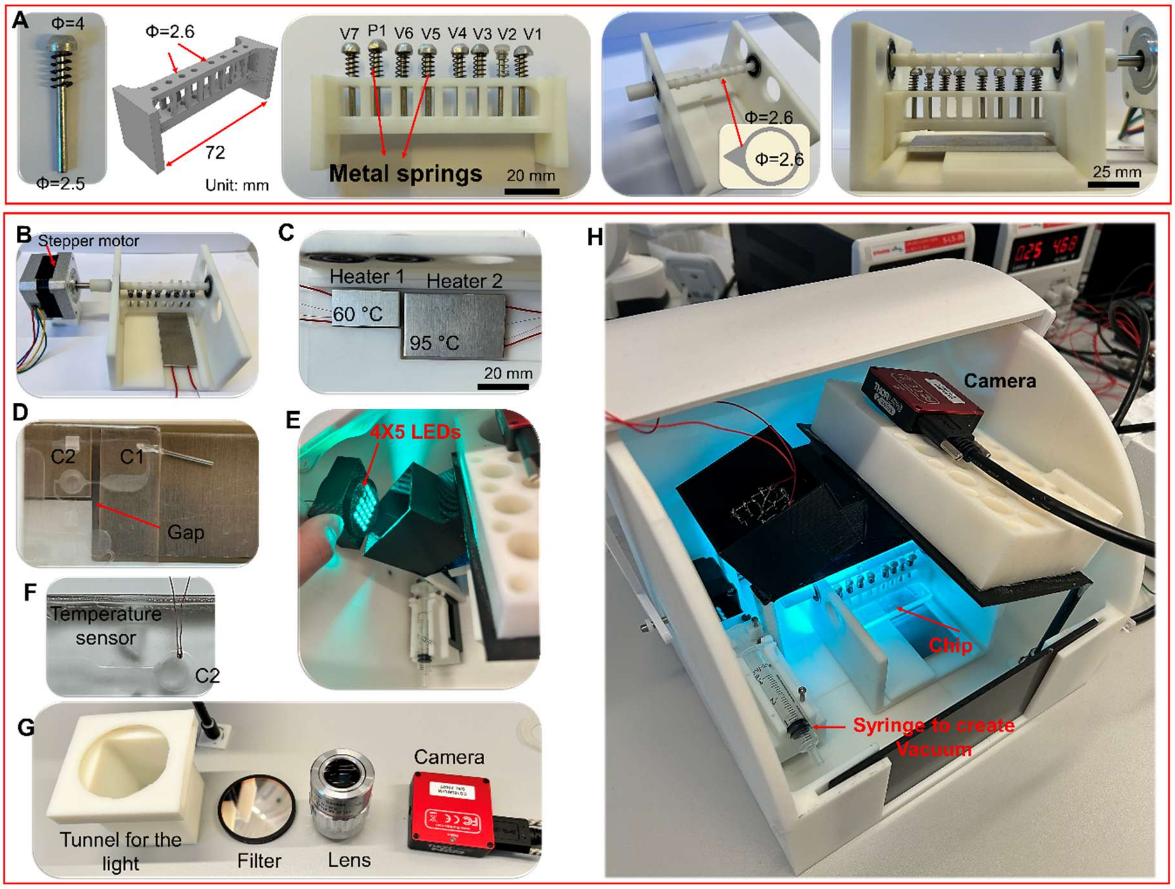


**Figure S1A. Total system for the multiplex diagnostic by real-time solid-phase PCR array. A)** The Mechanical CAM system to control the motion of the valves (V1-V7) and the fluid motion (P1). **B)** A stepper motor with coded motion creates the rotational motion in the shaft. The rotational motion then is converted into linear motion in the pins (CAM and follower system). Springs brings the pin to its original potion once the CAM pass its peak (steep) structure. **C)** The two heaters with different temperatures. **D)** The chip position on the heaters. C1 and C2 are located on heater 1 and heater 2, respectively. A small air gap (with 2 mm length) between the heaters are designed to prevent thermal gradient. **E)** The excitation light source (4x5 LEDs). **F)** Temperature sensor is inserted inside the chip to measure the temperature for thermal characterization of the PCR process. **G)** The disassembled optical system. **H)** The total box for the assay system.

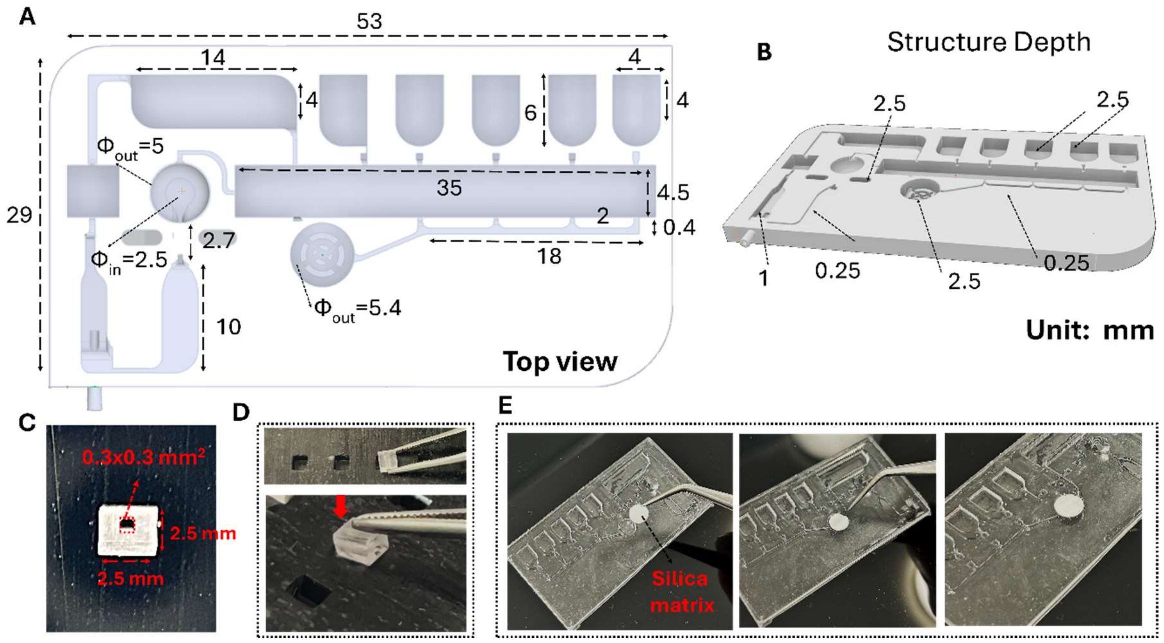


**Figure S1B. The chip dimension and the silica matrix insertion. A)** top view for the chip dimension in mm. **B)** The side view of the chip showing the depth of the channels and chambers. **C)** The rigid valve dimension. **D)** The valve insertion, with a slight push, in the flexible top layer. **E)** The silica matrix ($\Phi_{out}$=5.5mm) integrates into a 5.4 mm hole.

### CAM system

The CAM system is a mechanical motion transmission system where there is single rotational motion i. e. a stepper motor, is converted into a translational motion in the pins to press the valve and C2 (Figure S1 A and B).

### Fluid and temperature control

A mini-stepper motor, stepper 1 (T6 Mini Lead Screw Motion Stepper, ToAuto Tool) with small syringe (1ml) is placed beside the system to provide a vacuum in the chip to drive the fluids. sections to control the movement of the external magnet. Another stepper motor (stepper 2) is connected to the CAM system to rotate the followers, hence pressing the valve accordingly. Two customized heaters were placed under the device with a gap of 2 mm to prevent gradient temperature. The heaters were made from a 330 µm diameter nichrome wire (NI80-012, OMEGA Engineering inc.). for each heater, the wire was warped around a 3D-printed substrate that is attached to a customized aluminum block. A temperature sensor was placed in the middle of the aluminum block. The heating system was controlled by a heater controller (TEC-1089-SV, Meerstetter Engineering GmbH). An Arduino microcontroller (Arduino Uno R3, Arduino) controlled the total motions of the stepper motors and LEDs activation by a coded program in an open-source software (IDE, Arduino). To characterize the mixing efficiency in the mixing region, digital images were captured and analyzed using the Octave software. The mixing efficiency was calculated based on Equation S1.

$$\eta = \frac{\frac{1}{N}\sum_{i=1}^{N}(I_i - I_{0i})}{\frac{1}{N}\sum_{i=1}^{N}(I_r - I_{0i})} \times 100\% ,$$

where $I$ is the normalized gray scale value, $I$i is the value of each pixel at the ith pixel of the captured image, $I$r is the normalized value of the reference image in the fully mixed state, N is the number of pixels, and $I$0i is the normalized value at the $i$th pixel in the initial state, i.e., without mixing. Thus, we obtain η = 0% for the pre-mixed condition and η = 100% for the completely mixed condition.

The pressure generated inside the microchannel was measured using a pressure sensor (PX309–G5V, Omega). The total system components are listed in Table S1.

### Fluorescence measurements and image acquisition and processing

For optical detection systems, a fluorescent light source and detection are mounted on the top of the device. To cover a wide area on the chip and obtain strong excitation, the light source was built to have an array of 4x5 light-emitting diodes (LEDs) with a wavelength of 495 nm (HLMP-CE34-Y1CDD, Broadcom Inc.), a 495 ± 10 nm wavelength bandpass filter (35-878, Edmund Optics Ltd), and an optical achromatic lens (49-291, Edmund Optics Ltd) to focus the light source on the chip. The detection channel had a 510 ± 10 nm wavelength bandpass filter (49-291, Edmund Optics Ltd) and a monochrome CMOS Camera (CS165MU/M, Thorlabs). The fluorescence intensities of array spots are measured at the end of the annealing step. For each hybridization measurement, the average fluorescent intensity of 3 spots of the same capture target sequence was used. Initially, background adjustment was performed by subtracting average background fluorescence in the area of the array spots. Ct-values were considered by determining the cycle number at which the fluorescence intensity exceeded the standard deviation of average background intensity of 1-20 cycles. A grey bright-field image using a CMOS camera mounted on the top of the device was taken every cycle. Images were recorded at 10,000 frames/s with the exposure time of 1 or 2 µs. The captured images were transferred to the ThorCam software for analysis. The intensity of spots was extracted as a pixel peek value and plotted in real-time to determine the Ct values of the RT-PCR.

#### Array printing

To immobilize several probes, we tried the traditional array printing, wherein a microdroplet is simply placed on the top of the surface. However, we observed that for a relatively large spot area (~0.1 $mm^2$ ), a stronger probe appears in the side area of the spots (Shown in Figure 6). We hypothesize that the coffee-ring effect, where the drying process of the solute-laden droplet leads to the undesirable formation of a ring-like pattern of the solidified solute i. e. the probes, cause the undesirable distribution of the probes. Because probe immobilization requires several hours of solution incubation, drying the droplet is inevitable. Thus, we created a 3D-printed despising column array to immobilize the probe without drying (Figure S6). The despising array enables continuous fluid contact between the surface of the chip and the fluid (that contains the probe). The despising column works as an infinite liquid pool that supplies a probe from the top. Each column has a capacity of 2 µL. Using the despising column resulted in a uniform distribution of probe immobilization (See Figure S6). The suggested despising column also provides an easy and cheap tool with array up-scalability characteristic. (Figure S6) shows that the despising column array prints an array of 3 x 2) of spots. It is worth mentioning that the printing was performed inside a closed box at a maintained 98% relative humidity maintained in the printing enclosure. After the probe immobilization finished, the arrays were washed. The device was stored at room temperature in the vacuum-sealed bags until final integration.

**PCR efficiency and Assay sensitivity:**

To calculate the PCR efficiency, we run the RT-PCR with 10-fold dilution (starting with 10,000 copies) of the RNA. Then standard curve of the Ct-value for the real time PCR against the RNA target concentration is plotted. The slope of the standard curve is substituted in the PCR efficiency equation, $\boldsymbol{E} = (10^{(-1/\text{Slope})} - 1)$ X100. The assay sensitivity is evaluated by determining the assay ability to reliably detect the smallest quantity of a target nucleic acid i. e. limit of detection (LOD). There are several standard methods to assess PCR assay sensitivity, including serial dilution of target nucleic acid, standard curve analysis and replicates testing for statistical sensitivity. The serial dilution, where the lowest concentration at which the target is consistently detected (e.g., in 90% of tests) across multiple replicates, may suffer from several drawbacks of misassumption of dilution linearity, batch-to-batch variability and dilution factor variability i. e. the actual versus intended dilution factor. Hence, we used replicates testing as our statistical sensitivity assessment of the lowest detectable concentrations (1 copy/reaction). Spiked-in RNA with 1, 10, 100 and no-template controls (NTC) are repeatedly tested (6 times) to calculate the dilution level at which the assay detects the target in 95% of replicates. This ensures a robust determination of sensitivity.

**Table S1: Details of the chip components and their cost:**

| *Section* | *Component* | *Model* | *Company* | *Price (USD)* |
|---|---|---|---|---|
| Permanent parts | Stepper motor 1 | T6 Mini Lead Screw | ToAuto Tool | 52 |
| | Stepper motor 2 | NEMA17 | ToAuto Tool | 35 |
| | Controller | Arduino Uno R3 | Arduino | 12 |
| | Lens | 49-291 | Edmund Optics | 130 |
| | Excitation filter | 55-213 | Edmund Optics | 280 |
| | Emission filter | 35-878 | Edmund Optics | 280 |
| | Light-emitting diode | HLMP-CE34-Y1CDD | Broadcom Inc | 10 |
| | CMOS Camera | CS165MU/M | Thorlabs | 450 |
|  | Heater | --------- | Customized | 5 |
| | Heater Controller | TEC-1089-SV | Meerstetter Eng. | 300 |
|  | CAM system | --------------- | Customized | 5 |
| *Disposable parts* | 3D-printed valve | Customized | Customized | 0.1 |
| | 3D-Printed Mold for PDMS | Asiga Pico 2 | Asiga | 0.2 |
| | PDMS Chip | Sylgard 184 silicone | Diatom | 0.5 |

**Table S2. Primers for SARS-CoV-2 RPA**

Forward and reverse primers were designed to specifically identify the RNA of the following viruses

| Target virus | Primer name | Sequence 5'-3' | Target gene | PCR product size* |
|---|---|---|---|---|
| SARS-CoV-2 | SCoV_F | **FAM**-GATCTCAATGGTAACTGGTATGATTTCGGTG | RNA-dependent RNA polymerase | 119bp |
| | SCoV-R | GCCCTGGTCAAGGTTAATATAGGCATTAAC | | |
| | SCoV-P | **C6**-GAATCTACAACAGGAACTCCACTACCT | | - |
| Influenza A H1N1 | InfA-F | **FAM**-TAACGGGAAACTATGCAAACTAAGAGG | Hemagglutinin | 137bp |
| | InfA-R | GTGTTTCCACAATGTAGGACCATG | | |
| | InfA-P | **C6**-CTGTGGAGAGTGATTCACACTCTGGA | | - |
| Influenza B | InfB-F | **FAM**-GGGATAGAGATGGTACACGATGGTG | Neuraminidase | 105bp |
| | InfB-R | TGTGACAGTGTCCCATAGCAA | | |
| | InfB-P | **C6**-GGCTGTTGCAGCTGAATGCCAAGT | | - |
| Parainfluenza 1 | Pinf-F | **FAM-**TTCTGGAGATGTCCCGTAGG | | 204bp |
| | Pinf-R | CACATCCTTGAGTGATTAAG | | |
| | Pinf-P | **C6-**TTGCATCACCAATTGATAATGAAGG | | |
| Rhinovirus 89 | Rhin-F | **FAM-**AAGCACTTCTGTTTCCCCGG | | 208bp |
| | Rhin-R | CAGGCAGCCACGCAGGCTGG | | |
| | Rhin-P | **C6-**TCCAGCCTCATCTGCCAGGTCTACT | | |
| Immobilized FAM-probe | Immo-P | **C6-**GTAATTGATTAGCTTGTCGTTGTGA-**FAM** | - | - |

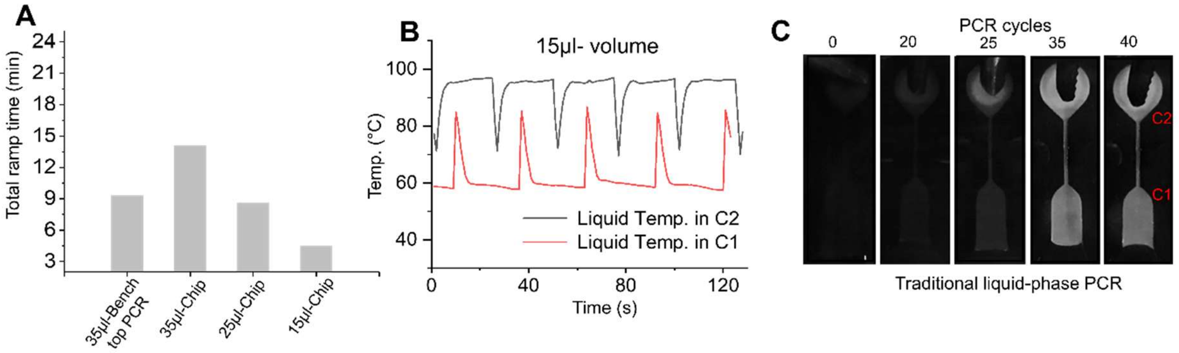


**Figure S2.** Thermal performance of the dual-chamber PCR system and liquid PCR. A) The total ramping time (heating and cooling) of the chip compared to commercial benchtop PCR cycler at different fluid volumes. B) The temperature inside the chambers wherein a sudden drop/rise temperature takes place the movement PCR mixture enters the specified chamber. C) Real images of traditional liquid-phase PCR over 40 PCR cycles.

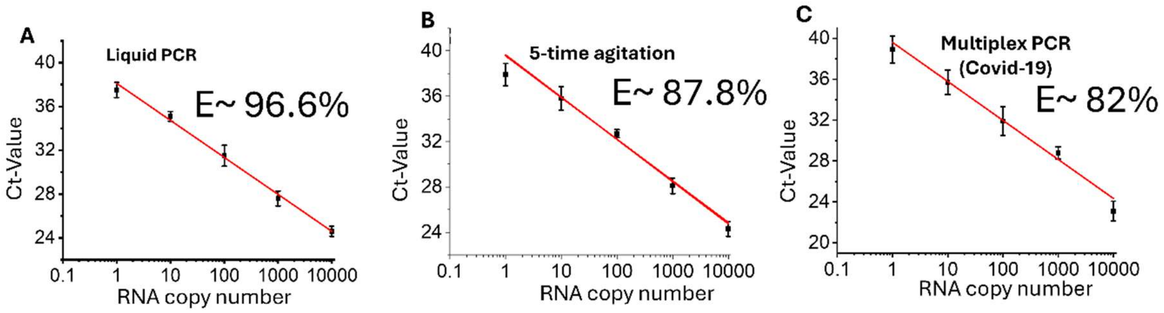


**Figure S3. The standard curve for PCR efficiency (E) calculation for : A**) Traditional liquid-phase PCR with CyberGreen dye. **B)** Solid-phase PCR with agitation during the annealing stage. **C)** Solid-Phase PCR with a multiple target existing in the chamber (data for Covid-19 virus).

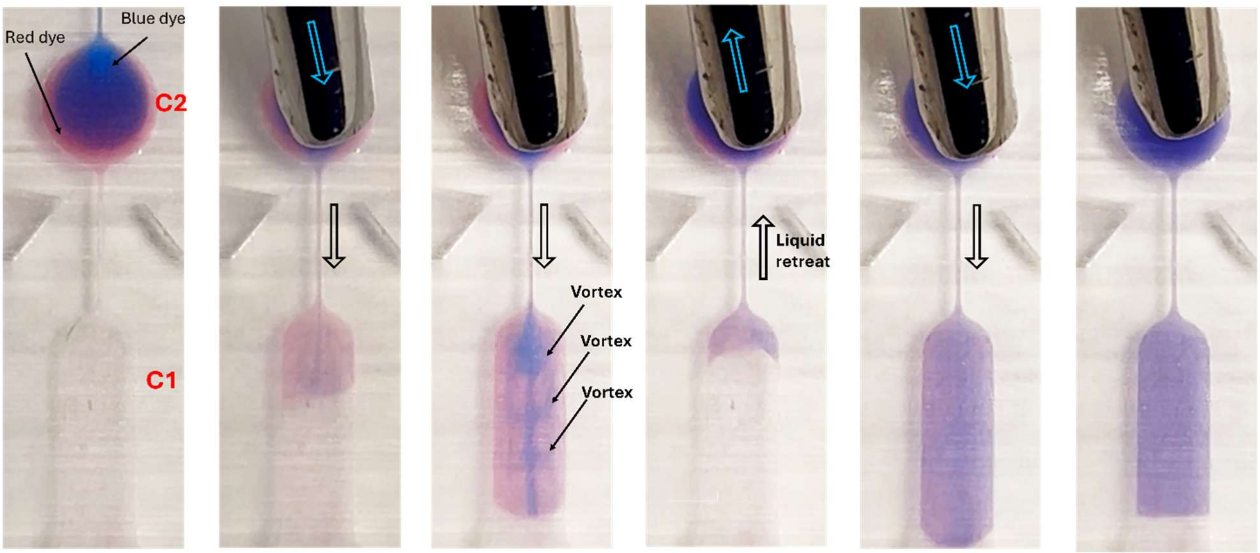


**Figure S4. Vortex creation during the fluid motion between C1 and C2.**

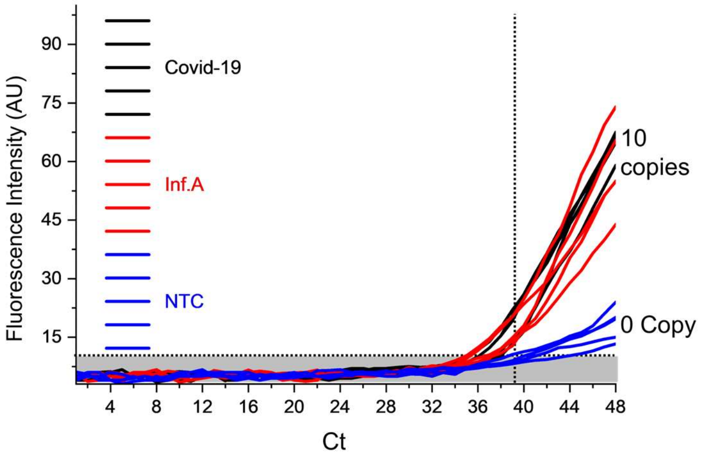


**Figure S5. Effect of late cycling.** The fluorescent intensity starts to increase after 39 cycles with non-template control (NTC) making the result of late PCR cycles unreliable for providing true positive due to non-specific amplification or primer-dimer formation.

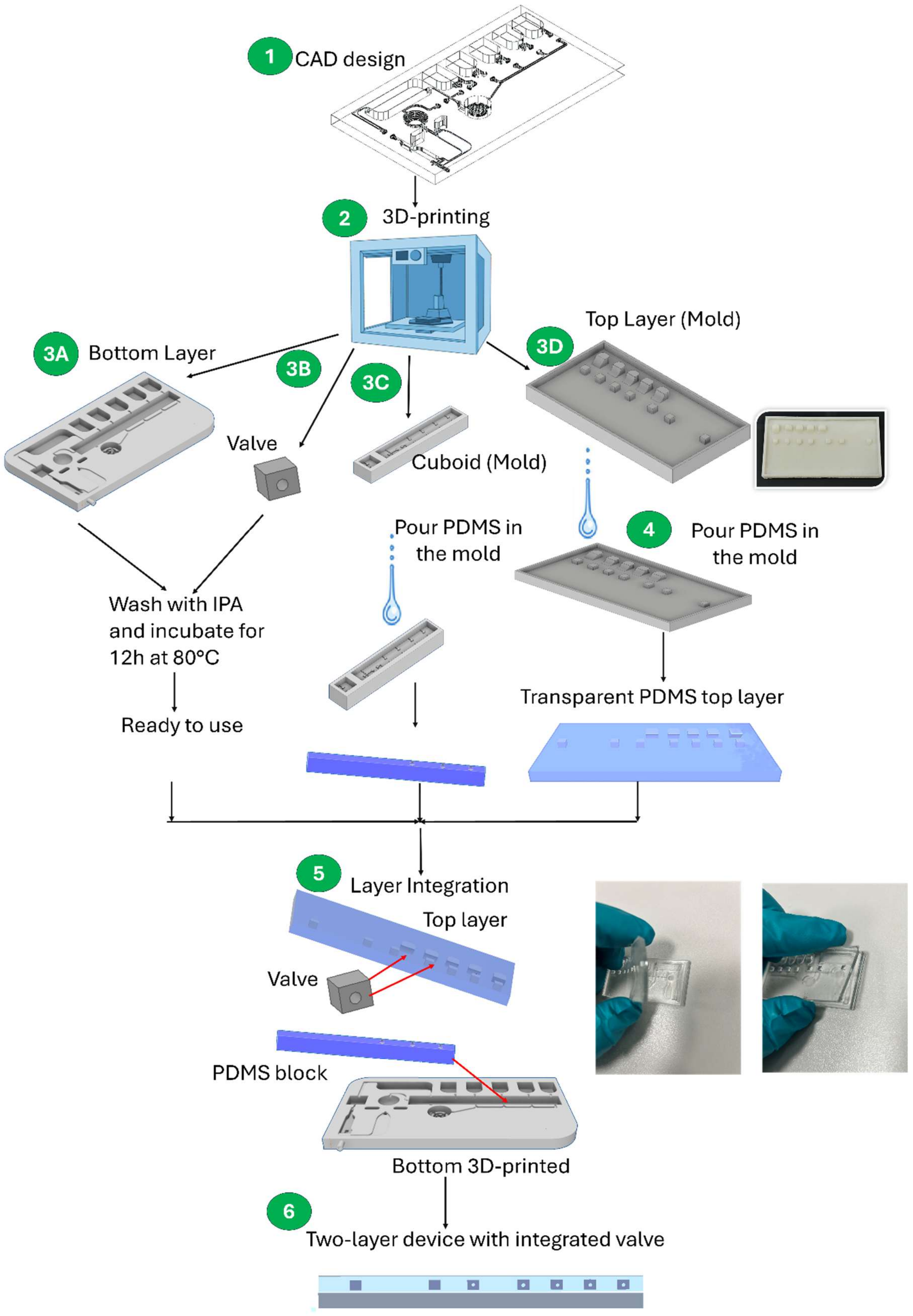


**Figure S6. Flow of the chip fabrication.** 3D-printer is used to print the bottom layer, the valve and the molds for top layer and the cuboid groove. The bottom layer and the valve parts are ready after printing. PDMS is poured into the molds to form the top layer and the cuboid shape, respectively. The cuboidal PDMS block is inserted physically inside the bottom layer while the valve is inserted physically into the top layer. To bond the 3D-printed bottom layer and PDMS bottom layer together, first the bottom layer is inserted into oxygen plasma machine (30 s) for surface activation to introduces polar functional groups (hydroxyl groups, –OH) onto the surface. Then, the bottom layer is submerged into APTES (silane coupling agent) to enable the triethoxysilane end reacting with hydroxyl groups on surface while the other side, amino group (-$NH_2$), is available for further chemical reaction. Finally, both top and bottom layers are inserted into oxygen plasma machine (30 s) prior to the final bonding of both layers.

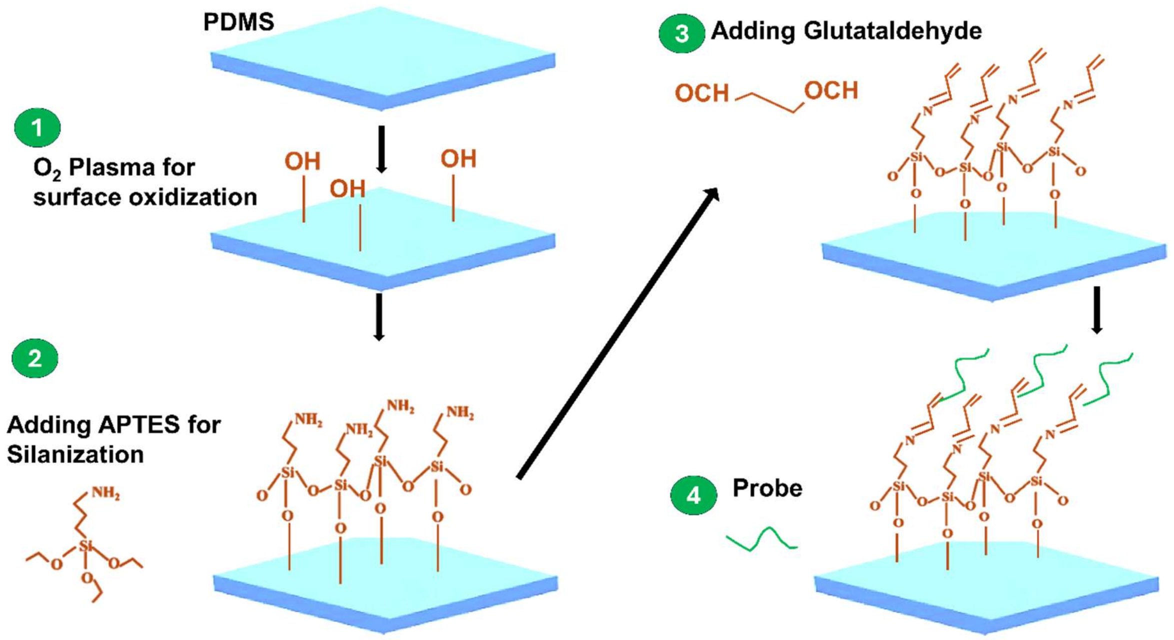


**Figure S7. Surface functionalization and probe immobilization**

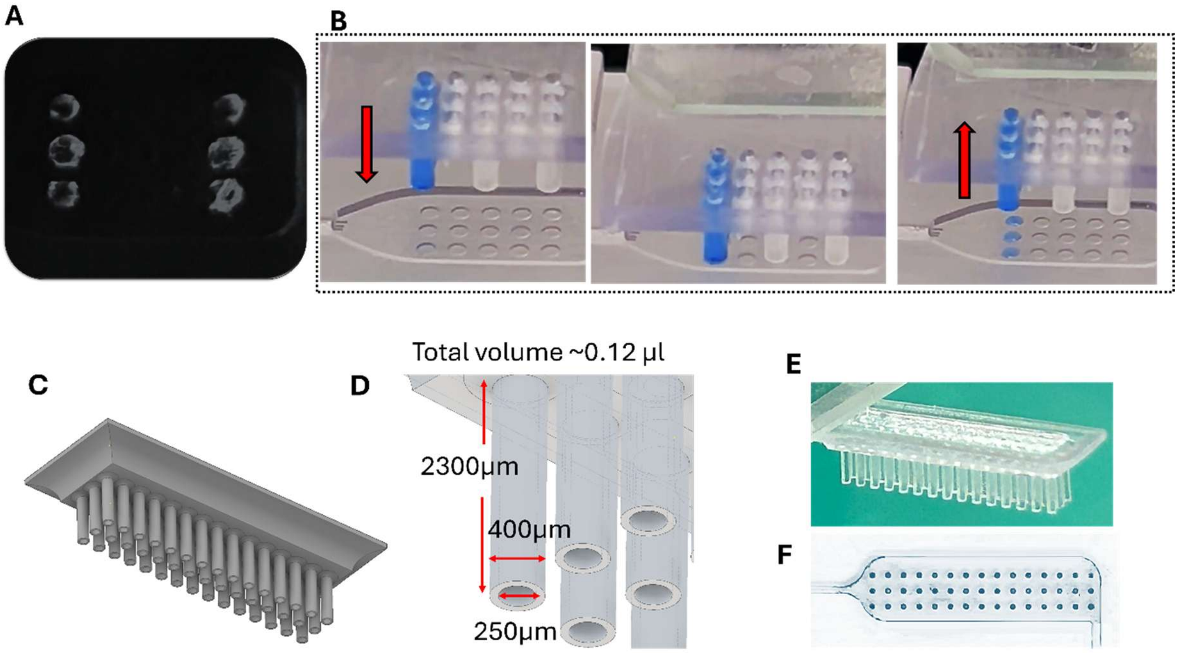


**Figure S8. Dispensing system for the array probe immobilization. A)** Non uniform fluorescence reading from the immobilized FAM due to, probably, the coffee ring effect, that occurs when a droplet containing particles dries on a surface, leaving a ring-like deposit of those particles or dried surface. **B)** Employing the column array dispenser developed by our team. **C-F)** CAD design, dimension and 3D-ringing of a possible scalable dispensing system for 45 spots (an array of 15X3).
